# *kri-1/KRIT1* restrains *skn-1/NRF2* activation to promote innate immune and lipid homeostasis

**DOI:** 10.64898/2026.05.15.725342

**Authors:** Akshay Alaghatta, Samantha Y. Tse-Kang, Daniel Bollen, Khursheed A. Wani, Nicholas D. Peterson, Read Pukkila-Worley

## Abstract

Animals must differentially allocate essential metabolic resources in response to changing environmental conditions to ensure reproductive success. For example, adult *C. elegans* coordinate innate immune defenses during infection and also allocate fat stores to the germline, each of which are required for evolutionary fitness. The genetic mechanisms that integrate resource allocation and host defenses in this context, however, are not fully understood. From a forward genetic screen for novel regulators of innate immune gene transcription, we identified *kri-1*, the nematode homolog of human Krev interaction trapped protein *KRIT1*. Mutations in human *KRIT1* underlie cerebral cavernous malformations, a disease marked by defects in vascular integrity. Similarly, *C. elegans kri-1/KRIT1* is required for the integrity of the intestinal epithelial cell barrier. We find also that *kri-1/KRIT1* is required for host tolerance to bacterial infection, fecundity, and normal development. In this context, *kri-1/KRIT1* restrains the activity of the cytoprotective transcription factor *skn-1/NRF2* to control immune gene transcription and intestinal lipid mobilization during aging, a process necessary for healthy reproduction, but functions independently of *skn-1*/*NRF2* to promote epithelial integrity and pathogen tolerance. These data reveal a broad role for a strongly conserved regulator in essential physiological processes that are required for reproductive fidelity and evolutionary success.

## INTRODUCTION

Evolutionary success for an individual organism is defined by its ability to survive to reproductive age and produce progeny. In this context, animals must defend against environmental threats and coordinate the metabolic resources that are necessary to support germline function. The genetic pathways that control these essential physiological processes are therefore required for organismal evolution.

In their natural habitats, the microscopic nematode *C. elegans* ensures its evolutionary fidelity by navigating biotic and abiotic threats to produce large broods [1, 2]. One nematode, for example, produces 250-300 progeny to ensure successful passage of its genome to the next generation. Also of note, *C. elegans* eat bacteria as their source of nutrition and can be infected with pathogens in the wild [3]. As a result, intestinal epithelial cells evolved mechanisms to defend against ingested pathogens, which are essential for nematodes to survive to reproductive age [1, 2, 4–8]. Therefore, *C. elegans* provides a facile experimental platform to characterize the genetic architecture that evolved to integrate stress responses, pathogen resistance mechanisms, and resource allocation with the shared goal of ensuring reproductive success for individual animals.

Reproduction requires extensive mobilization of metabolic resources in all animals. In *C. elegans*, lipids from the soma are shuttled to the germline [9–12]. Somatic depletion of fat compromises overall longevity of the animal, but ensures reproductive fitness by supporting gonadal function [13]. For example, germline-less *C. elegans* animals have increased somatic lipid stores, which enhance longevity-promoting cellular programs, such as proteostasis and perhaps also immunity [11, 12]. Coordination of this energy balance is complex, with multiple regulators operating in different tissues, which are responsive to changing environmental conditions. One such regulator that is particularly important in this context is the conserved cytoprotective transcription factor SKN-1, homologous to mammalian NRF2 [14–17]. Constitutive activation of SKN-1 drives lipids to the germline, which supports normal fecundity, but at the expense of stress resistance and survival in nutrient poor conditions [15].

Many of the key insights regarding how *C. elegans* survives environmental threats to reach reproductive age has come from forward genetic screens. For example, the *C. elegans* p38 PMK- 1 immune pathway was uncovered in a screen for mutants that rendered animals hypersusceptible to bacterial infection – a seminal moment for the field of *C. elegans* innate immunity, as it demonstrated that nematodes coordinate inducible anti-pathogen defenses through evolutionarily conserved pathways [4]. Our group and others subsequently used forward genetics to reveal mechanisms of pathogen sensing and immune regulation [9, 10, 18–22]. One approach that has been useful in this context is use of *gfp*-based transcriptional reporters for innate immune effector genes, screening for mutations that cause constitutive activation of *gfp*. A forward genetic screen for regulators of an immune reporter for infection response gene (*irg*)-5 *(irg-5*p*::gfp)*, for example, led to the discovery that the p38 PMK-1 pathway is activated during pathogen infection by a regulator that is expressed on lysosome-related organelles [18, 19]. A screen with a different reporter (*irg-4*p*::gfp*) identified a sensory neuron-to-intestine axis that suppresses p38 PMK-1 innate immune activation to promote growth and development [20].

Here, we report the results of a forward genetic screen to characterize the regulation of the immune effector *irg-6*. We discover that mutations in *C. elegans kri-1*, the nematode homolog of human Krev interaction trapped 1 (*KRIT1*), cause constitutive activation of *irg-6* and other immune effector genes. Accordingly, *kri-1* mutants have compromised host tolerance to infection, slower rates of development and smaller brood sizes, phenotypes that are each rescued by re-introduction of wild-type *kri-1* in the *kri-1* mutant strain. In addition, *kri-1* mutants have defects in the integrity of the intestinal epithelial barrier, an intriguing finding in light of the fact that mutations in human KRIT1 cause defects in vascular permeability that cause cerebral cavernous malformations. To investigate the mechanism by which *kri-1* regulates key physiological processes that are required for evolutionary fitness, we examined the transcriptome of *kri-1* mutants and observed that *kri-1* suppresses the transcription of genes that are targets of the cytoprotective transcription factor SKN-1/NRF2. We find that *kri-1* controls the nuclear translocation of SKN-1 to regulate the mobilization of lipids from the soma to the germline, a process required for reproduction. Intriguingly, KRIT1 is required for the nuclear localization of NRF2 in mammalian cells, suggesting that this mechanism is strongly conserved through animal evolution [23, 24]. However, we found that *kri-1* functions independently of *skn-1* to regulate pathogen tolerance, epithelial barrier integrity, and developmental rate. These data uncover a conserved regulator that links lipid metabolism, epithelial barrier integrity, and host defense to ensure the evolutionary success of an individual animal.

## RESULTS

### Forward genetic screen identifies *kri-1*/*KRIT1* as a suppressor of innate immune gene transcription

We conducted a forward genetic screen to identify new regulators of the innate immune response in *C. elegans*. We previously identified three innate immune effectors genes (*irg-4*, *irg-5*, and *irg-6*) that are each strongly induced during infection and required to promote pathogen resistance – RNAi-mediated knockdown of *irg-4, irg-5,* and *irg-6* each render *C. elegans* hypersusceptible to killing by *P. aeruginosa* [20, 25–27]. The basal expression (*i.e.*, in the absence of pathogen infection) of *irg-4* and *irg-5* requires the p38 PMK-1 innate immune pathway [19, 20]. Previously, we performed forward genetic screens using *gfp*-based transcriptional reporters for *irg-4* and *irg-5* that identified regulators of the p38 PMK-1 pathway. Unlike *irg-4* and *irg-5*, the basal expression of *irg-6* is not dependent on the p38 PMK-1 pathway [6]. In addition, *irg-6* was the single most strongly induced gene in wild-type *C. elegans* during *P. aeruginosa* infection in two different whole genome transcription profiling experiments [5, 26]. Therefore, we sought to understand how the innate immune effector gene *irg-6* is regulated as a means to identify p38 PMK-1-independent mechanisms of immune regulation in *C. elegans*.

As in our previous forward genetic screens, we generated a transcriptional immune reporter for *irg-6* by fusing its promoter to *gfp* and expressing it from an extrachromosomal array. We screened the F2 progeny from ∼29,000 mutagenized haploid genomes and identified fifteen mutants that caused constitutive *irg-6*p*::gfp* expression (**Fig. 1A**). To identify the causative mutations, we performed whole-genome sequencing of F2 recombinants that were homozygous for constitutive *irg-6*p*::gfp* expression (pooled from 3X or 4X backcrosses to the parent *irg-6*p*::gfp* strain). Six of the 15 mutant alleles contained putative loss-of-function mutations in the gene *kri-1*: five mutants (*ums26, ums27, ums28, ums29*, and *ums31*) had a nonsense mutation (Trp336*) and another allele (*ums21*) had a splice donor mutation (**Fig. 1B**). KRI-1 is the nematode homolog of human KRIT-1 and has three protein domains (NUDIX-5, Ankyrin repeats and a FERM B-lobe). The mutations in the *kri-1* alleles recovered from this screen were found in or before the second of these domains (**Fig. 1B**).

**Figure 1.**
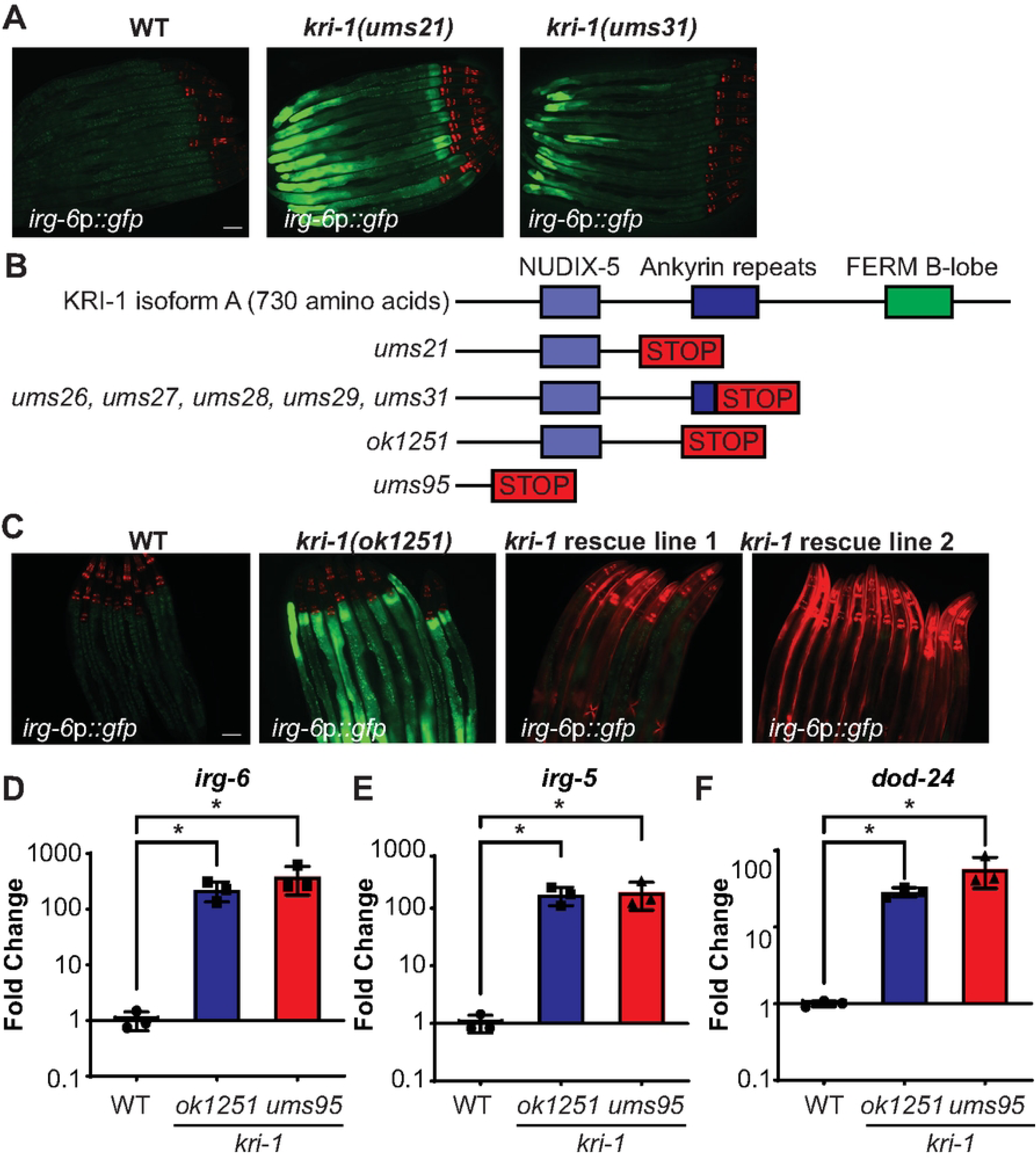
Forward genetic screen identifies *kri-1/KRIT1* as a suppressor of innate immune gene transcription. **A.** Images of *C. elegans irg-6*p*::gfp* transcriptional reporter expression in wild-type animals and in two *kri-1* mutant strains isolated from the forward genetic screen. **B.** Diagram of *kri-1* mutants. **C.** Images of *C. elegans irg-6*p*::gfp* transcriptional reporter expression in wild-type animals, *kri-1(ok1251)* mutants, and two independent *kri-1* transgenic rescue strains [*kri-1(ok1251) + kri-1*]. **D.** qRT-PCR of the indicated genes. Data are the average of three independent replicates, each normalized to a control gene, with error bars representing SEM. Data are presented as the value relative to the average expression from all replicates of the indicated gene in wild-type animals. *p < 0.05 (one-way ANOVA) for the indicated comparison.

Several lines of evidence indicate that loss-of-function mutations in *kri-1* cause constitutive activation of innate immune effector genes. We found that the previously characterized, loss-of-function allele *kri-1(ok1251)* also caused constitutive *irg-6*p::GFP activation (**Fig. 1C)** [28]. Importantly, expression of *kri-1* under its endogenous promoter in the *kri-1(ok1251)* mutant background from two independent transgenic lines restored wild-type expression of *irg-6p*::GFP (**Fig. 1C**). Using RT-qPCR, we tested whether mutation of *kri-1* increased expression of the native *irg-6* gene, as well as other immune effectors (**Fig. 1D-F**). *C. elegans kri-1(ok1251)* mutants constitutively express *irg-6* (**Fig. 1D**), *irg-5* (**Fig. 1E**), and *dod-24* (**Fig. 1F**). Finally, we used CRISPR-Cas9 to disrupt the start codon of *kri-1* to create an additional loss-of-function strain, *kri-1(ums95)*. The immune effectors *irg-5, irg-6,* and *dod-24* were upregulated in *kri-1(ums95)* mutants to similar levels as observed in the *kri-1(ok1251)* background (**Fig. 1D-F**). Thus, *kri-1* suppresses innate immune gene expression in *C. elegans*.

### Innate immune dysregulation in *kri-1* mutants compromises host tolerance to infection

To examine the broader impact of *kri-1* on immune gene transcription, we analyzed a published dataset of genes that are misregulated in *kri-1(ok1251)* mutants [29]. We performed a gene set enrichment analysis (GSEA) to compare *kri-1*-regulated genes with the group of genes that are differentially transcribed in wild-type *C. elegans* infected with *P. aeruginosa* [30]. Consistent with the qRT-PCR data of *irg-6*, *irg-5,* and *dod-24* (**Fig. 1D-F**), genes that are normally induced during *P. aeruginosa* infection are enriched among the genes that are upregulated in *kri-1(ok1251)* mutants (**Fig. 2A**). Thus, *kri-1* suppresses the transcription of pathogen-response genes.

**Figure 2.**
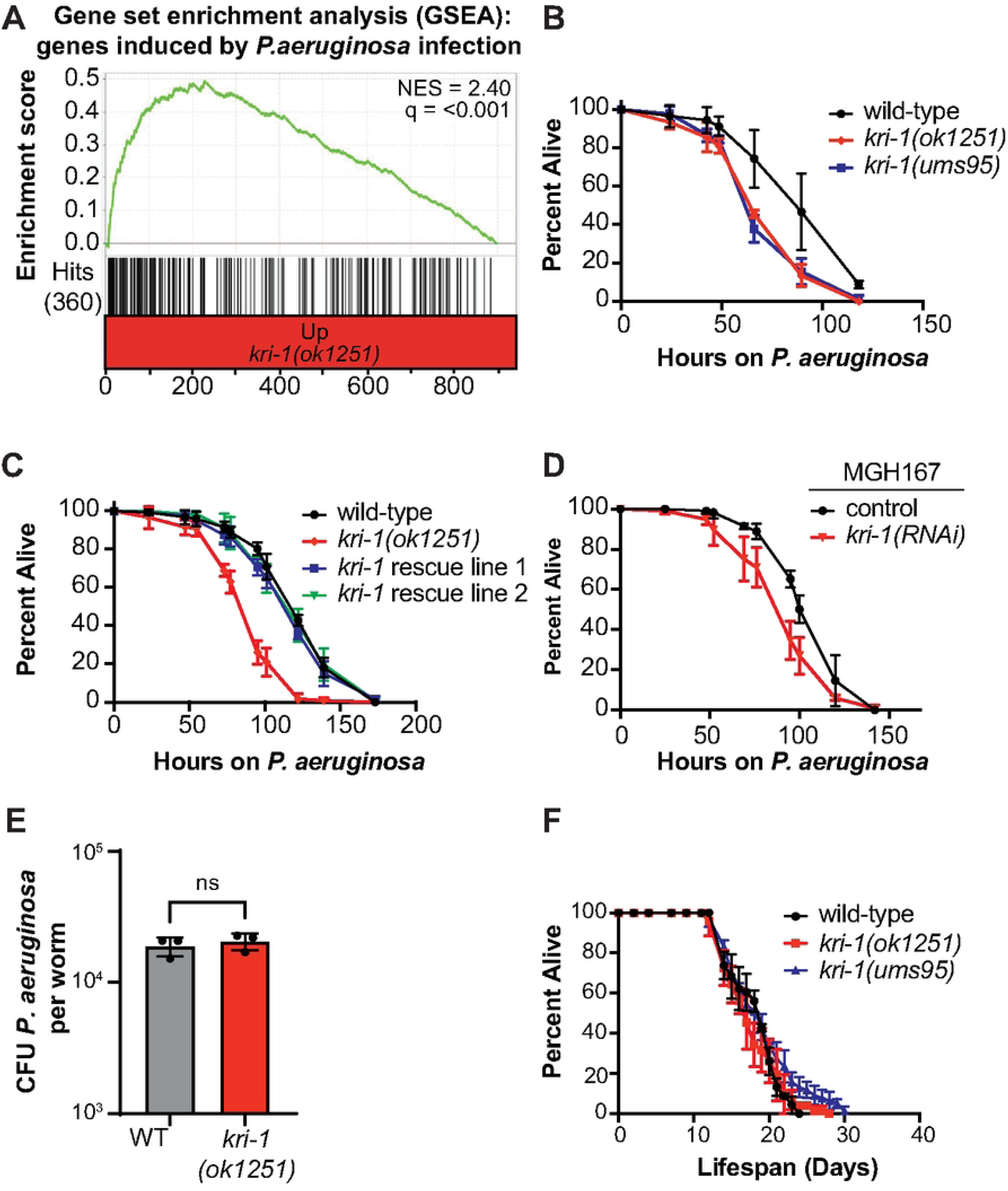
Innate immune dysregulation in *kri-1* mutants compromises host tolerance to infection. **A.** Gene set enrichment analysis (GSEA) of genes upregulated during infection in mRNA-seq experiment comparing *kri-1(ok1251)* mutants with wild-type animals. **B, C and D.** *P. aeruginosa* pathogenesis assays with the indicated *C. elegans* genotypes. In C, *C. elegans* strain MGH167 was used, which permits RNAi only in intestinal cells. For each of the assays in B, C, and D, Data are the average of three replicates with error bars indicating SEM. Two to three trials of each assay was performed. p<0.05 (log rank) for difference between the *kri-1* mutants or *kri-1(RNAi)* and the other genotypes. Sample sizes, mean lifespan, and p-values for each trial are shown in Table S1. **E.** Colony-forming units (CFUs) of *P. aeruginosa* in the intestine after 48 h of bacterial infection in *C. elegans* with the indicated genotypes. Data are the average of three replicates (n = 10 animal per replicate). Source data is in Table S1. **F.** The lifespans of wild-type, *kri-1(ok1251)* and *kri-1(ums95)* mutant animals are shown. Data are the average of four replicates with error bars indicating SEM. Three trials of each assay was performed. p = not signifcant (log rank) for difference between the genotypes. Sample sizes, mean lifespan, and p-values for all trials are shown in Table S1.

We found that *kri-1(ok1251) and kri-1(ums95)* loss-of-function mutants are hypersusceptible to killing by *P. aeruginosa* (**Fig. 2B**), a finding that has also been observed in mutations of other immune regulators that cause aberrant immune activation [19, 21, 27, 31]. Re-introduction of *kri-1* expressed under its own promoter from two independent extrachromosomal arrays restored wild-type resistance to *P. aeruginosa* infection in the *kri-1(ok1251)* mutant (**Fig. 2C**). In addition, knockdown of *kri-1* only in the intestine, using a transgenic *C. elegans* strain engineered only to perform RNAi just in this tissue, enhanced pathogen-induced mortality (**Fig. 2D**). These data suggest that *kri-1*, which is expressed in the intestinal epithelium [28], acts cell-autonomously within intestinal cells to promote pathogen resistance. We quantified the colony-forming units (CFUs) of *P. aeruginosa* in the intestine of *kri-1(ok1251)* mutants during infection. Interestingly, at 48 hours of infection, intestinal CFUs of *P. aeruginosa* in *kri-1(ok1251)* mutants were similar as in wild-type animals (**Fig. 2E**). These data indicate that the immune dysregulation in *kri-1* mutants compromises host tolerance to infection, rather than disrupting the anti-pathogen response. Importantly, two *kri-1* loss-of-function mutants did not shorten the normal lifespan of *C. elegans* (**Fig. 2F**). Thus, *kri-1* functions to support host tolerance of infection without pleiotropic effects on nematode lifespan.

### *kri-1/KRIT1* supports intestinal barrier integrity and evolutionary fitness

KRI-1 is a cytoskeletal protein that is expressed in the *C. elegans* intestine. Notably, human KRIT1 supports the integrity of the epithelial barrier, and mutations in *KRIT1* contribute to cerebral cavernous malformations (CCM) disease, which is characterized by leakage of blood due to increased permeability [32, 33]. We therefore determined if *kri-1* supports the integrity of the intestinal barrier in *C. elegans.* We used an intestinal dye that allows quantification of intestinal barrier permeability by assessing leakage of blue “Smurf” dye from the intestinal lumen into the body cavity of *C. elegans*. We observed that *C. elegans kri-1(ok1251)* mutant animals accumulated more blue dye in the soma than wild-type worms (**Fig 3A-B**). Re-introduction of *kri-1* expressed under its own promoter suppressed the leakage phenotype of the *kri-1(ok1251)* mutant (**Fig. 3A-B**). We corroborated this observation using a GFP-based translational reporter for ACT-5 (ACT-5::GFP), the isoform of actin that is expressed in the intestine [34]. ACT-5 forms the apical terminal web, which supports the microvillar brush border and provides cytoskeletal support for epithelial junctions. RNAi-mediated knockdown of *kri-1* disrupted the apical actin cytoskeleton in intestinal epithelial cells compared to control animals (**Fig. 3C-D**).

**Figure 3.**
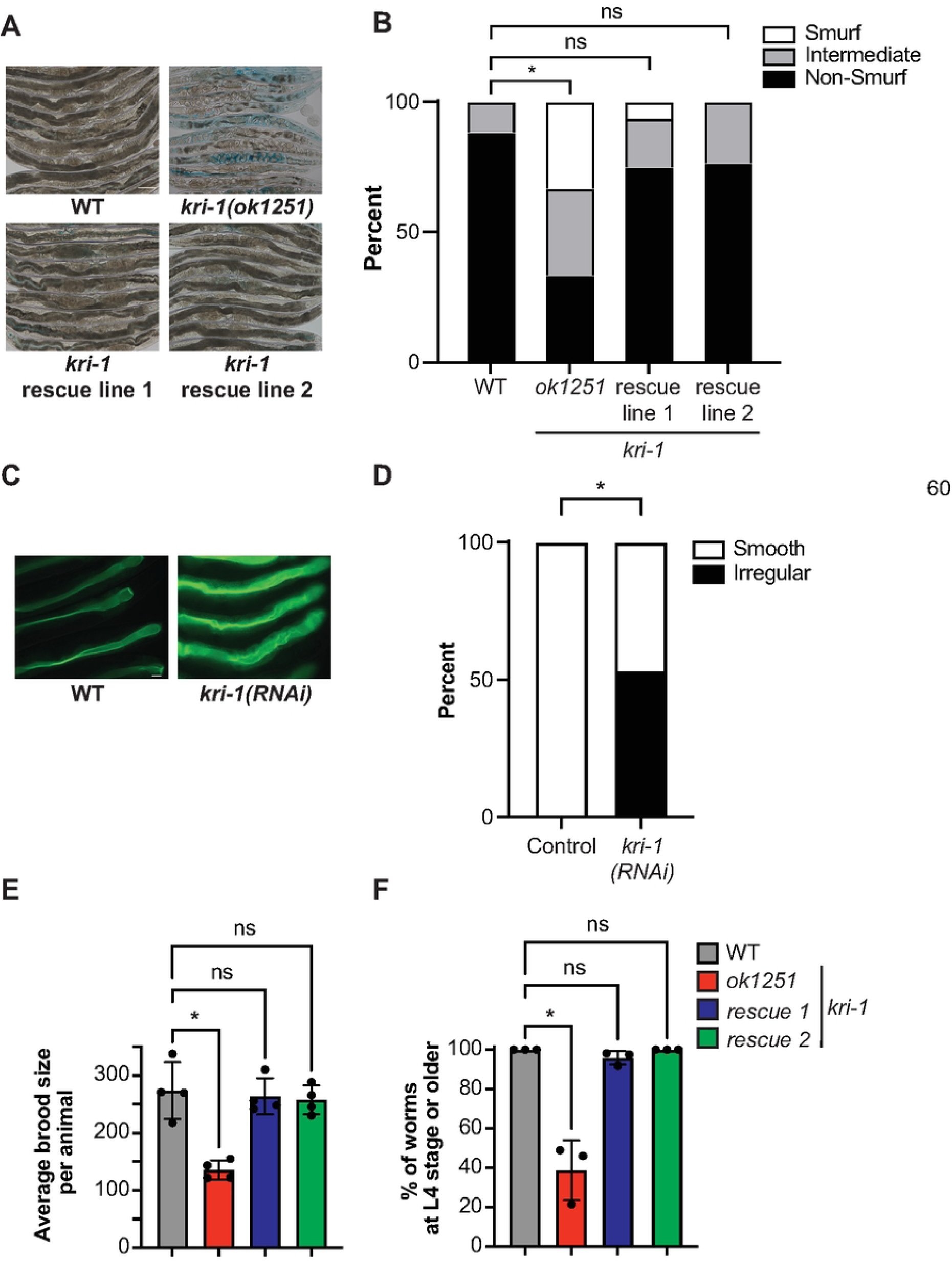
*kri-1/KRIT1* supports intestinal barrier integrity and evolutionary fitness A and. **B.** *C. elegans* animals of the indicated genotypes were examined using the “smurf” assay, which assesses the permeability of the intestinal barrier. Percentage of animals exhibiting “non-smurf”, “intermediate”, or “smurf” phenotype in a qualitative assessment is shown. * p<0.05 (Fisher’s exact test). Scale bar: 100 μM. Sample sizes and p-values are shown in Table S1. **C and D.** *C. elelgans* ACT-5::GFP animals were subjected to control or *kri-1(RNAi).* Data are presented as the percentage of animals exhibiting “smooth” or “irregular” luminal structure. * p<0.05 (Fisher’s exact test) for the indicated comparison. Scale bar: 20 μM. The data for each replicate are presented in Table S1. **E.** Brood sizes from animals of the indicated genotypes were quantified. Each data point is the average brood size from three or four animals. * p < 0.05 (one-way ANOVA). The data for each replicate are presented in Table S1. **F.** Development assays were performed with the indicated genotypes. The stage of the animals was recorded approximately 72 h after egg lay. Data are presented as the average percent of animals for each genotype that were L4 or older (percent L4+) from three replicates, with error bars representing SEM. * p < 0.05 (one-way ANOVA).

We also observed that *kri-1(ok1251)* mutants are slow growing and have smaller brood sizes than wild-type controls. We quantified the development of *C. elegans* to the L4 stage in *kri-1(ok1251)* mutants, as well as in the two *kri-1* rescue lines in which expression of this gene was restored in the mutant background (**Fig. 3E**). We observed that developmental timing was comparable to wild-type in the two *kri-1* rescue lines. Similarly, the brood size of *kri-1(ok1251)* mutants was significantly smaller than wild-type animals and was also restored to wild-type levels in the two *kri-1* rescue lines (**Fig. 3F**).

Together, these data demonstrate that *C. elegans kri-1* is required for both intestinal barrier integrity and evolutionary fitness.

### KRI-1/KRIT1 restrains SKN-1/NRF2 activity to control intestinal lipid mobilization during aging

*C. elegans kri-1* functions with the cytoprotective transcription factor SKN-1/NRF2 to promote lifespan extension in germline-less animals, which have slower rates of aging and are used to study genetic mechanisms that extend lifespan [28, 35]. In this context, *kri-1* regulates the redox species H_2_S and reactive oxygen species (ROS) production, which activate SKN-1 [28, 35]. Consistent with these data, we found that SKN-1 targets were enriched among the genes hyperactivated in *kri-1(ok1251)* mutants (**Fig. 4A**). We performed a GSEA on genes that were previously identified as SKN-1 targets in an RNA-seq experiment of the *skn-1(lax188)* gain-of-function mutant (**Fig. 4A**). Genes are that negatively regulated, as well as those that are positively regulated, by SKN-1, were enriched among genes that were upregulated and downregulated, respectively, in *kri-1(ok1251)* mutants compared to wild-type controls (**Fig. 4A**). These data suggest that *kri-1* suppresses SKN-1 activation. Of note, the group of genes that are both activated by SKN-1 and suppressed by KRI-1 includes a significant enrichment of pathogen-response genes (**Fig. 4B**).

**Figure 4.**
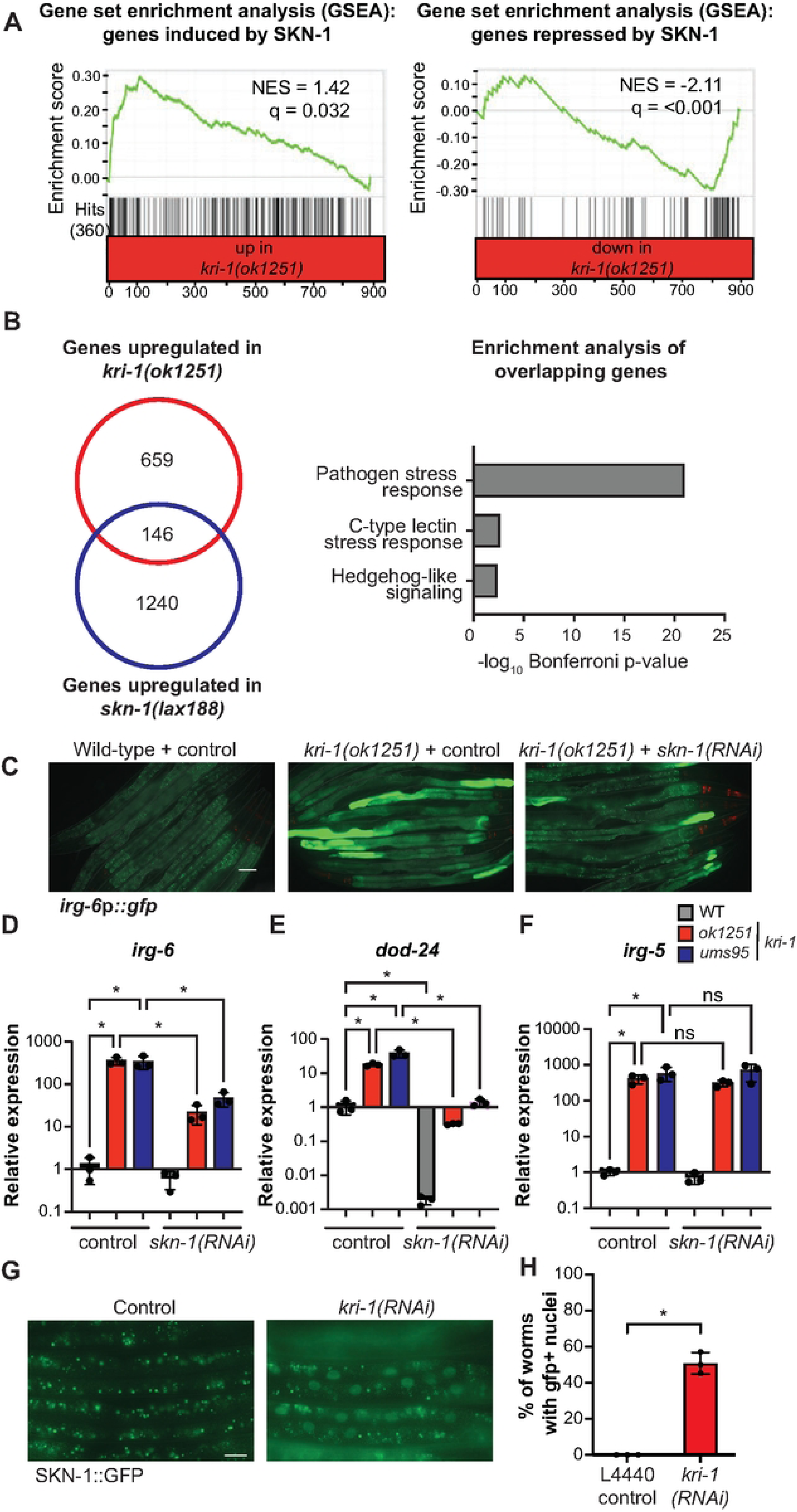
KRI-1/KRIT1 suppresses SKN-1/NRF2. **A.** Gene set enrichment analyses (GSEA), which compare the genes that are differentially regulated in *skn-1(lax188)* gain-of-function and *kri-1(ok1251)* loss-of-function mutants, with the comparisons indicated in the left and right panels. **B.** Overlap of genes differentially regulated in the indicated comparisons and an enrichment analysis of overlapping genes. **C.** Images of *irg-6*p*::gfp* expression in the indicated genoytpes. **D, E and F.** qRT-PCR of the indicated genes. Data are the average of three independent replicates, each normalized to a control gene, with error bars representing SEM. Data are presented as the value relative to the average expression from all replicates of the indicated gene in wild-type animals. *p < 0.05 (one-way ANOVA) for the indicated comparison. **G and H.** Images and quanification of accumulation of SKN-1::GFP in the nuclei of intestinal epithelial cells in the indicated genotypes. Data are presented as the average percent of animals that exhibited SKN-1::GFP nuclear localization, with error bars representing SEM. *p < 0.05 (one-way ANOVA). Source data is in Table S1.

We performed gene expression experiments to characterize the regulation of immune effector genes by KRI-1 and SKN-1. Consistent with the GSEA data, RNAi-mediated knockdown of *skn-1* suppressed the hyperactivation of *irg-6*p*::gfp* in the *kri-1(ok1251)* mutant background (**Fig. 4C**). A qRT-PCR analysis of the native *irg-6* locus (**Fig. 4D**) and *dod-24* (**Fig. 4E**) confirmed this observation in two separate *kri-1* loss-of-function mutant backgrounds (*ok1251* and *ums95*). However, the regulation of *irg-5* by *kri-1* occurs independently of *skn-1* (**Fig. 4F**). Upon activation, SKN-1 translocates from the cytoplasm into the nucleus, which can be quantified in a transgenic *C. elegans* strain in which the SKN-1 protein was tagged with GFP. Consistent with KRI-1’s role as a negative regulator of SKN-1, RNAi-mediated knockdown of *kri-1* caused SKN-1 protein to differentially accumulate in the nucleus (**Fig. 4G-H**). Thus, KRI-1 inhibits SKN-1 activation.

To determine the physiological relevance of KRI-1-mediated inhibition of SKN-1 activation, we first studied *C. elegans* infection with *P. aeruginosa.* Unlike *kri-1* mutants, the *skn-1(lax188)* gain-of-function mutants were not hypersusceptible to killing by *P. aeruginosa* in a model of intestinal infection called the “slow kill” assay (**Fig. 5A**). We previously found that *skn-1(lax188)* gain-of-function mutants are resistant to *P. aeruginosa* in the “fast kill” assay, a different model of pseudomonal pathogenesis that assesses the toxicity of secreted phenazine toxins. However, *kri-1(ok1251)* mutants demonstrated wild-type susceptibility to phenazine intoxication in this assay (**Fig. 5B**). Similarly, *skn-1(lax188)* gain-of-function mutants did not have reduced brood size (**Fig. 5C**). We also found that RNAi-medicated knockdown of *skn-1* did not suppress the defect in intestinal barrier integrity of the kri-1(ok1251) mutants, as quantified using the Smurf dye, described above (**Fig. S1A-B**). Thus, KRI-1 does not function with SKN-1 to support pathogen tolerance, fecundity, or epithelial barrier integrity.

**Figure 5.**
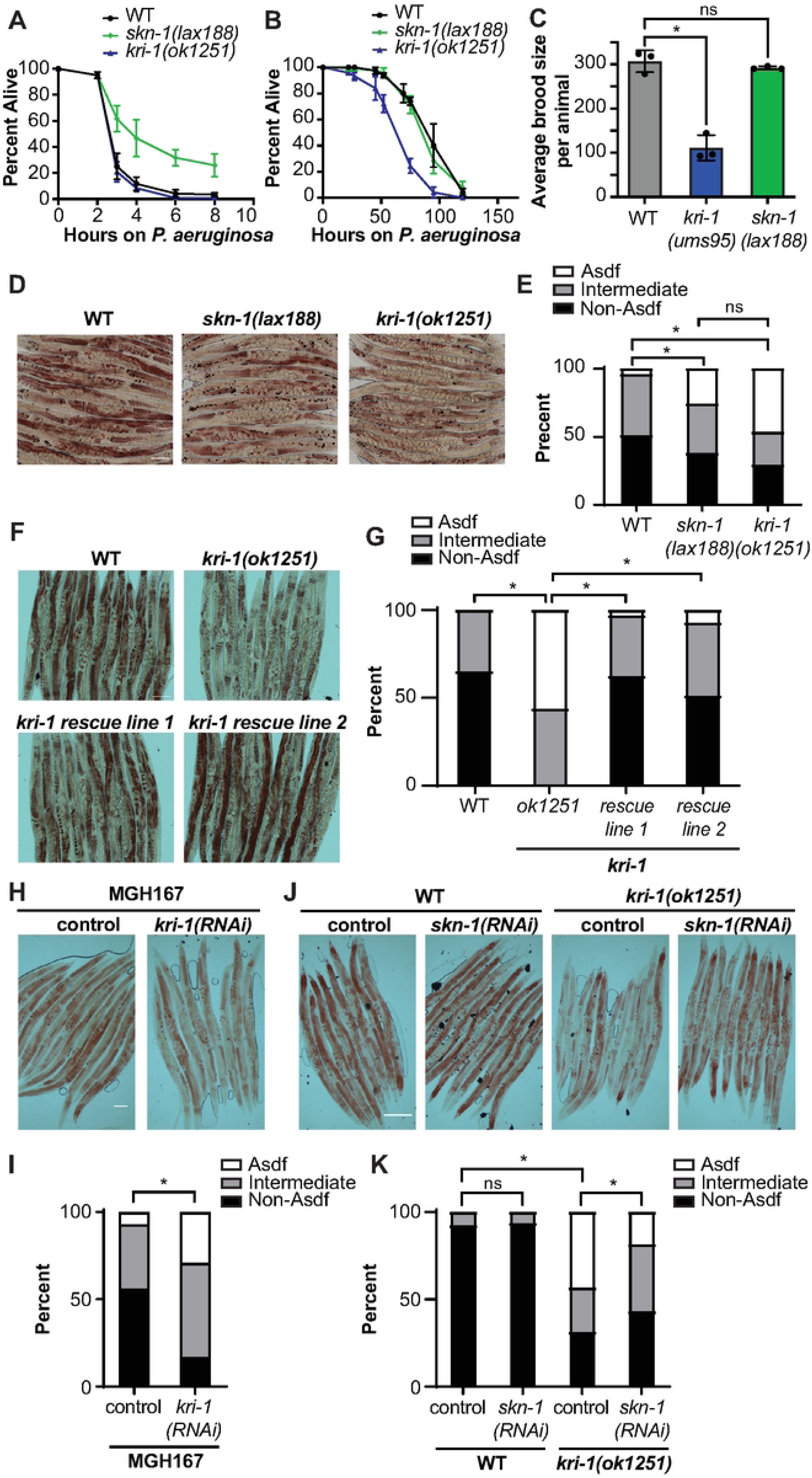
KRI-1/KRIT1 restrains SKN-1/NRF2 activity to control intestinal lipid mobilization during aging. Representative *P. aeruginosa* pathogenesis assays in C. elegans. **A.** “fast-kill” assay. **B.** “slow kill” assay. Data are the average of three biological replicates with error bars representing SEM. Sample sizes, mean lifespan, and p-values for each trial are shown in Table S1. **C.** Brood sizes from animals of the indicated genotypes were quantified. Each data point is the average brood size from three animals. * p < 0.05 (one-way ANOVA). The data for each replicate are presented in Table S1. **D, E, F, G, H, I, J and K.** Oil-red-O staining of *C. elegans* animals of the indicated genotypes. Data are the average number of animals displaying Asdf, intermediate or non-Asdf phenotypes. *C. elegans* strain MGH167 permits RNAi only in intestinal cells. * p <0.05 (Fisher’s exact test). Sample sizes and p-values are shown in **Table S1**.

In contrast, we found that SKN-1 and KRI-1 cooperate to promote adaptive changes in lipid metabolism during aging. SKN-1 drives the redistribution of somatic lipids from intestinal epithelial cells to the germline to ensure reproductive and evolutionary fitness as *C. elegans* age [14–17]. Constitutive activation of SKN-1 in the *skn-1(lax188)* gain-of-function allele causes marked depletion of intestinal lipid stores in Day 2 adults – a phenotype called age-dependent depletion of somatic fat (ASDF) that can be quantified using the lipophilic dye Oil-red-O [14–17]. Intriguingly, aging *kri-1(ok1251)* animals display the ASDF phenotype similar to the *skn-1(lax188)* allele, consistent with our findings that SKN-1 is constitutively activated in the *kri-1* mutant background (**Fig. 5D-E**). Re-introduction of *kri-1* expressed under its own promoter in two independent rescue lines complemented the excessive depletion of somatic fat observed in the *kri-1(ok1251)* mutant (**Fig. 5F-G**). In addition, RNAi-mediated knockdown of *kri-1* only in intestinal tissues also caused excessive depletion of somatic fat, indicating that, like *skn-1*, *kri-1* functions cell-autonomously in this context as well (**Fig. 5H-I**). Importantly, RNAi-mediated knockdown of *skn-1* suppressed the ASDF phenotype of *kri-1(ok1251)* mutants (**Fig. 5J-K**). Thus, KRI-1 restrains SKN-1/NRF2 to control intestinal lipid mobilization to the germline during aging.

## DISCUSSION

Here, we characterize a key role for *kri-1/KRIT1* in the regulation of innate immune effector expression, lipid metabolism, and intestinal barrier integrity, which is required to support reproductive fidelity and evolutionary fitness in *C. elegans*. From a forward genetic screen for regulators of immune effector transcription, we uncovered *kri-1,* the nematode homolog of human *KRIT1*. We show that *kri-1* is required for host tolerance to infection and restrains *skn-1*/*NRF2* activation to control intestinal lipid mobilization during aging. *C. elegans* KRI-1 suppressed the transcription of genes that are induced by SKN-1, as well as the translocation of activated SKN-1 to the nucleus. Importantly, *kri-1* loss-of-function mutants phenocopied *skn-1* gain-of-function mutants with respect to immune effector induction and excessive mobilization of lipids from the soma to the germline. RNAi-mediated knockdown of *skn-1* in the *kri-1* mutant background suppressed these phenotypes, establishing the physiological relevance of this genetic interaction in the context of reproduction. We also find that *kri-1* functions independently of *skn-1* to regulate intestinal barrier integrity and pathogen tolerance, suggesting that KRI-1 may function upstream of other cellular regulators in the control of key physiological processes that are required for *C. elegans* to reach reproductive age.

Mutation of *KRIT1*, the human orthologue of *C. elegans kri-1,* underlies cerebral cavernous malformations (CCM), a condition in which compromised vascular integrity leads to blood leakage and the formation of numerous vascular lesions in the brain [32, 33]. In this context, *KRIT1* functions to maintain endothelial tight junctions in capillaries and may also be required for immune signaling and intestinal microbial homeostasis [36, 37]. Similiarly, we found that loss of *C. elegans kri-1* disrupts the organization of the intestinal actin isoform ACT-5 and compromises intestinal epithelial barrier integrity, phenotypes that are reminiscent of the dysregulated filamentous actin observed in mammalian endothelial cells following loss of KRIT1 [38]. In addition, KRIT1 is required for the nuclear localization of NRF2 in mammalian endothelial cells, fibroblasts, and hepatic cells [23, 24]. The direct parallels between the studies of mammalian *KRIT1* and *C. elegans kri-1* suggest that the mechanism of *kri-1*/*KRIT1* in the regulation of essential physiological processes is strongly conserved through animal evolution.

These data also demonstrate that a *kri-1* - *skn-1* axis is required for lipid homeostasis and protective mobilization of lipids from the soma to the germline. NRF2 is a central regulator of lipid mobilization in mammals [23, 24]. Of relevance, NRF activation in KRIT1-deficient hepatic cells perturbs metabolic pathways under stress conditions [24]. These data raise the intriguing possibility that the role of KRIT1 in the regulation of lipid metabolism is also strongly conserved. The mechanism by which *C. elegans kri-1* inhibits *skn-1* activation is not known, however. One possibility is that KRI-1 binds to SKN-1 in the cytoplasm and physically restrains it from entering the nucleus. Alternatively, damage to epithelial integrity in the *kri-1* mutants could be an upstream activator of a cytoprotective response regulated by SKN-1.

In summary, we identify a single regulator that is strongly conserved across the animal branch of the tree of life and show that it is a lynchpin coordinator of essential host responses, which are required for reproductive fidelity and evolutionary success of an animal.

## ACKNOWLEDGMENTS

The authors thank Sean Curran and Alex Soukas for helpful discussions, and Melanie Trombly for critical reading of the manuscript. This research was supported by R01 AI130289 (to R.P.W.), R01 AI159159 (to R.P.W.), F30 DK127690 (to S.Y.T.), T32 AI095213 (to S.Y.T.), F30 AI150127 (to N.D.P.), and T32 AI132152 (to N.D.P.). Some strains were provided by the *Caenorhabditis* Genetics Center, which is funded by the NIH Office of Research Infrastructure Programs (P40 OD010440). The funders had no role in study design, data collection and analysis, decision to publish, or preparation of the manuscript.

## MATERIALS AND METHODS

### C. elegans strains

The previously published *C. elegans* strains used in this study are: N2 Bristol (wild type), CF2052 *kri-1(ok1251)* [28], SPC227 *skn-1(lax188)* [39], SPC2002 SKN-1::GFP [16], MGH167 *sid-1(qt9);alxIs9*[*VHA-6p::SID-1::SL2::GFP*] [40].

The strains reported in this study are: AU308 *agIs41[irg-6p::GFP; myo-2p::mCherry]*, RPW325 *kri-1(ums21); agIs41[irg-6p::GFP; myo-2p::mCherry]*, RPW326 *kri-1(ums24); agIs41[irg-6p::GFP; myo-2p::mCherry]*, RPW327 *kri-1(ums26); agIs41[irg-6p::GFP; myo-2p::mCherry]*, RPW328 *kri-1(ums27); agIs41[irg-6p::GFP; myo-2p::mCherry]*, RPW329 *kri-1(ums28); agIs41[irg-6p::GFP; myo-2p::mCherry]*, RPW330 *kri-1(ums29); agIs41[irg-6p::GFP; myo-2p::mCherry]*, RPW331 *kri-1(ums31)*; *agIs41*[*irg-6*p*::GFP; myo-2*p*::mCherry*], RPW548 *kri-1(ok1251)*; *agIs41*[*irg-6*p*::GFP; myo-2*p*::mCherry*], RPW549 *kri-1(ums95)*, RPW580 *kri-1(ok1251)*; *umsEx139*[*kri-1*p*::kri-1(cDNA); myo-3*p*::mCherry*]; *agIs41*[*irg-6*p*::GFP; myo-2*p*::mCherry*] *(Line 1)*, RPW582 *kri-1(ok1251)*; *umsEx139*[*kri-1*p*::kri-1(cDNA); myo-3*p*::mCherry*]; *agIs41*[*irg-6*p*::GFP; myo-2*p*::mCherry*] *(Line 2)*.

*C. elegans* strains were maintained on nematode growth medium (NGM) plates [0.25% Bacto-peptone, 0.3% sodium chloride, 1.7% agar (Fisher), 5 μg/mL cholesterol, 25 mM potassium phosphate pH 6.0, 1 mM magnesium sulfate, 1 mM calcium chloride] with *E. coli* OP50 as a food source.

### Bacterial strains

Bacteria used in this study were *Escherichia coli* OP50, *E. coli* HT115(DE3), and *Pseudomonas aeruginosa* strain PA14.

*E. coli* OP50 was grown in LB supplemented with 0.175 mg/mL streptomycin at 37 °C for 16–18 h at 250 rpm. *P. aeruginosa* strain PA14 was grown in LB at 37 °C for 14–15 h at 250 rpm.

### Forward genetic screen

Ethyl methanesulfonate (EMS) mutagenesis was performed on the AU308 *agIs41 (irg-6*p::*gfp)* strain as previously described [19–21]. Briefly, synchronized L4 animals were treated with 48.6 mM EMS in M9 buffer for 4h at 22 °C on a rotator. Mutagenized P0 animals were plated on NGM plates seeded with *E. coli* OP50. Gravid F1 progeny were treated with hypochlorite, and the resulting eggs were hatched overnight to obtain synchronized F2 animals. F2 progeny were divided into six distinct pools and screened for bright, constitutive *irg-6*p::*gfp* expression. Approximately 29,000 mutagenized haploid genomes were screened and fifteen mutants were identified.

To identify the causative allele, whole-genome sequencing of F2 recombinants that were homozygous for constitutive *irg-6*p*::gfp* expression, pooled from 3X or 4X backcrosses to the parent *irg-6*p*::gfp* strain, was performed. Genomic DNA was isolated using the Gentra Puregene DNA Isolation Kit (Qiagen) and submitted for whole-genome sequencing (BGI). Libraries were sequenced using 100 bp paired-end reads, yielding an average of 65 million reads per sample and ∼130× genome coverage. Homozygous variants relative to the WBcel235 (ce11) reference genome that were present in mutants, but absent from the *agIs41* parental strain were identified using in-house pipelines, as described [10, 19]. Homozygous variants were called using ‘bcftools’. Variants present in both the parent *agIs41* strain and the forward genetic mutants were removed using ‘bedtools’. Remaining homozygous variants were annotated with ‘snpEff’ using the *C. elegans* reference genome WBcel235.99.

### *C. elegans* strain generation

CRISPR–Cas9 editing with single-stranded oligodeoxynucleotide (ssODN) homology-directed repair was used to generate *kri-1(ums95)*, as previously described [18, 19, 26, 41]. All CRISPR reagents were purchased from Integrated DNA Technologies. Guide sequences were selected using the CHOPCHOP web tool [42]. The sequence for the ssODN repair template contained the indicated mutation flanked by 35 bp homology arms and is given in **Table S2**. F1 progeny were screened for Rol (roller) phenotypes 3–4 days after injection. PCR and Sanger sequencing was used to confirm the edit. Genotyping primer sequences are listed in **Table S2**.

To generate the *irg-6*p::*gfp* reporter strain, 1.0 kb of DNA upstream of the *irg-6* start codon was amplified from wild-type genomic DNA and fused to *gfp*, amplified from the pPD95.75 vector, using PCR. 10 ng/µL of the promoter::*gfp* fusion was microinjected into the gonads of young adult hermaphrodites along with 8 ng/µL *myo-3*p*::mCherry* co-injection marker and 100 ng/µL of 1-kb ladder DNA. The resulting extrachromosomal array was inherited by a subset of progeny, initially selected based on pharyngeal mCherry expression, and subsequently integrated into the genome by UV irradiation to create a stable transgenic line.

Generation of transgenic rescue strains was performed as described previously [18–20, 26, 27, 41]. The *kri-1* promoter (∼2 kb upstream of the ATG start codon), *kri-1* coding sequence, and *kri-1* 3′ UTR were amplified by PCR and cloned into a pUC19 vector using Gibson assembly. 5 ng/µL of this plasmid was microinjected along with 5 ng/µL of pCFJ104 (*myo-3p::mCherry*) and 100 ng/µL of empty pUC19 plasmid into strain AU308 *agIs41(irg-6p::GFP;myo-2p::mCherry)*, which was subsequently crossed with strain RPW548 *kri-1(ok1251); agIs41(irg-6*p::*GFP;myo-2p::mCherry)* to generate both rescue lines.

### *C. elegans* bacterial pathogenesis and lifespan assays

“Slow-killing” *P. aeruginosa* infection assays were performed as described [43, 44]. Briefly, *P. aeruginosa* PA14 was grown overnight and 10 µL of the culture was seeded onto the center of 35-mm tissue culture plates containing 4 mL of slow-kill agar (0.35% Bacto-peptone, 0.3% NaCl, 1.7% agar, 5 µg/mL cholesterol, 25 mM potassium phosphate, 1 mM MgSO_4_, 1 mM CaC1_2_). Plates were incubated for 24 hours at 37 °C and then for an additional 24 hours at 25 °C. L4-stage *C. elegans* animals were transferred onto PA14 slow-kill plates supplemented with 0.1 mg/mL 5-Fluoro-2’-deoxyuridine (FUDR) to prevent progeny production and maintained at 25 °C. Animals were scored for survival twice daily until the end of the experiment.

“Fast-killing” *P. aeruginosa* infection assays were performed as previously described [18, 26, 45]. In brief, a single colony of *P. aeruginosa* was inoculated into 3 mL LB Lennox medium and incubated at 37 °C for 14 h with shaking at 250 rpm. Five microliters of this culture was then spread in the center of 35-mm tissue culture plates containing 4 mL fast-kill agar (PSG agar: 1% Bacto-peptone, 1% glucose, 1% sodium chloride, 150 mM sorbitol, 1.7% Bacto-agar). Plates were incubated for 24 h at 37 °C followed by 24 h at 25 °C. Approximately 40 L4-stage nematodes were transferred to the pseudomonal lawns. Dead animals were scored at 2, 4, 8, or 24 h by assessing movement after tapping the head with a platinum wire.

*C. elegans* lifespan assays were conducted following an established protocol, with animals grown on nematode growth media agar seeded with *E. coli* OP50 at 20 °C in the presence of 0.1 mg/mL FUDR [43]. Animals were scored for survival daily until the end of the experiment.

At least three independent biological replicates of each assay were performed. Sample sizes, mean survival, and statistical comparisons for each trial are reported in **Table S1**.

### Oil Red O staining

Oil Red O (ORO) staining of lipids was performed as previously described with minor protocol modifications [14]. Briefly, Day 2 adult age-matched worms were collected in 1 mL of M9 buffer in a microcentrifuge tube and centrifuged at 25 × g for 1 minute. The supernatant was aspirated, and worms were washed with 1 mL of M9 buffer and centrifuged. The supernatant was again aspirated until ∼100 μL remained, followed by the addition of 600 μL of 60% isopropanol. Samples were rotated for 3 min at room temperature and centrifuged at 25 × g. The supernatant was aspirated again until ∼100 μL remained. Worms were then stained for 2 h with 3 mg/ml ORO solution while rotating at room temperature, followed by a final wash in M9 buffer before collection for brightfield imaging and quantification.

Quantification was conducted as previously described [14, 15]. Fat levels were subjectively categorized into three groups: non-Asdf, intermediate, and Asdf. Non-Asdf worms exhibit dark red ORO staining throughout most of the soma. Intermediate Asdf phenotype is depletion of somatic fat with some ORO staining in the soma present. Worms with the Asdf phenotype have near-complete loss of observable somatic fat deposits.

### Intestinal barrier function with Smurf assay

Smurf staining was performed as previously described with minor modifications [46]. To summarize, age-matched animals were incubated for 3 h in liquid cultures of *E. Coli* OP50 bacteria supplemented at a 1:1 ratio with blue food dye (Spectrum FD&C Blue #1 PD110, 5.0% wt/vol in water). Animals were then recovered and examined for the presence or absence of blue dye in the body cavity, described as a “smurf” phenotype, using brightfield microscopy. Worms were sorted as “smurf,” “intermediate,” or “non-smurf” and analyzed with a Fisher’s exact test using GraphPad Prism.

### Development, brood size, and CFU assays

Development assays were conducted following a previously established protocol [20, 43]. Briefly, a 2 hour egg lay was performed, and hatched animals were grown on *E. coli* OP50 for 72 hours at 20°C. Larval stages were then visually quantified under a dissecting microscope. Animals at the L4 stage or older were counted. Three biological replicates were performed.

Brood size assays were conducted following a previously established protocol [20, 43]. Briefly, brood sizes were quantified from 3-5 independent plates per condition, each with 2-3 animals per plate. Animals were transferred to new plates each day to facilitate scoring of the progeny.

Colony-forming units (CFUs) of *P. aeruginosa* within the *C. elegans* intestine were quantified as previously described, with minor modifications [43]. Briefly, animals were exposed to *P. aeruginosa* for 48 h and then transferred to NGM plates lacking bacteria for 10 min to allow removal of external bacteria. Animals were subsequently washed with M9 buffer containing 0.01% Triton X-100, and homogenized with a bead mill homogenizer. CFUs were quantified from serial dilutions of the resulting lysate.

Source data for all figures can be found in **Table S1**.

### Feeding RNAi

*C. elegans* were fed *E. coli* HT115 expressing dsRNA targeting the genes of interest. In brief, HT115 bacteria expressing the relevant dsRNA constructs were streaked onto LB agar plates containing 50 μg/mL ampicillin and 15 μg/mL tetracycline and incubated overnight at 37 °C. Individual colonies were then inoculated into LB broth with 50 μg/mL ampicillin and grown overnight at 37 °C for 16–18 h with shaking at 250 rpm. These overnight cultures were seeded onto NGM plates supplemented with 5 mM IPTG and 50 μg/mL carbenicillin and incubated for 16–18 h at 37 °C to induce dsRNA expression. Synchronized L1 stage animals were transferred onto the prepared RNAi plates and allowed to develop to the L4 stage.

### Microscopy and image analysis

For fluorescence and brightfield imaging, animals were mounted on 2% agarose pads, immobilized with 50 mM tetramisole (Sigma), and imaged using a Zeiss AXIO Imager Z2 microscope equipped with a Zeiss Axiocam 506 mono camera and Zen 2.5 (Zeiss) software.

### Quantification and statistical analysis

Differences in survival in *P. aeruginosa* pathogenesis or lifespan assays were assessed using log-rank tests on Kaplan–Meier survival curves. OASIS 2 was used for all survival analyses [47]. GSEA analyses were performed as described [48, 49]. WormCat 2.0 was used for gene enrichment analyses [50]. Statistical hypothesis testing for other experiments was performed in Prism 10 (GraphPad) using the methods indicated in the figure legends. **Table S1** contains all source data, as well as the sample sizes, survival data, and p-values for all trials of phenotypic assays reported in this paper.

## FIGURE LEGENDS

**Figure S1. *kri-1/KRIT1* functions independently of SKN-1 to support intestinal barrier integrity.** *C. elegans* animals of the indicated genotypes were examined using the “smurf” assay, which assesses the permeability of the intestinal barrier. Percentage of animals exhibiting “non-smurf”, “intermediate”, or “smurf” phenotype in a qualitative assessment is shown. * p<0.05 (Fisher’s exact test). Scale bar: 100 μM. Sample sizes and p-values are shown in Table S1.

**Table S1. Source Data, including sample sizes, statistical tests, and p values for all data in this manuscript.**

**Table S2. Primer, crRNA guide and ssODN sequences designed for this study**

